# Inactivation of medial frontal cortex changes risk preference

**DOI:** 10.1101/390021

**Authors:** Xiaomo Chen, Veit Stuphorn

**Affiliations:** Department of Neuroscience, Johns Hopkins University School of Medicine and Zanvyl Krieger Mind/Brain Institute, 3400 N. Charles St., Baltimore, MD 21218-2685, USA; Department of Psychological and Brain Sciences, Johns Hopkins University, 3400 N. Charles St., Baltimore, MD 21218-2685, USA; Current address: Department of Neurobiology, Stanford University School of Medicine, Stanford University, Clark Center W1.3, 318 Campus Drive West Stanford, CA 94305

## Abstract

Humans and other animals need to make decisions under varying degrees of uncertainty. These decisions are strongly influenced by an individual’s risk preference, however the neuronal circuitry by which risk preference shapes choice is still unclear [1]. Supplementary eye field (SEF), an oculomotor area within primate medial frontal cortex, is thought to be an essential part of the neuronal circuit underlying oculomotor decision-making, including decisions under risk [2–5]. Consistent with this view, risk-related action value and monitoring signals have been observed in SEF [6–8]. However, such activity has also been observed in other frontal areas, including orbitofrontal [9–11], cingulate [12–14], and dorsal lateral frontal cortex [15]. It is thus unknown whether the activity in SEF causally contributes to risky decisions, or if it is merely a reflection of neural processes in other cortical regions. Here, we tested a causal role of SEF in risky oculomotor choices. We found that SEF inactivation strongly reduced the frequency of risky choices. This reduction was largely due to a reduced attraction to reward uncertainty and high reward gain, but not due to changes in the subjective estimation of reward probability or average expected reward. Moreover, SEF inactivation also led to increased sensitivity to differences between expected and actual reward during free choice. Nevertheless, it did not affect adjustments of decisions based on reward history.

## Results

### Monkeys are risk-seeking

In our gambling task, two monkeys (*Macaca mulatta*, A and I) had to choose between two gambles with different combinations of maximum reward amount and winning probability (Figure 1A and Methods). Risk was quantified as reward uncertainty, using standard economic models [16–18].

**Figure 1.**
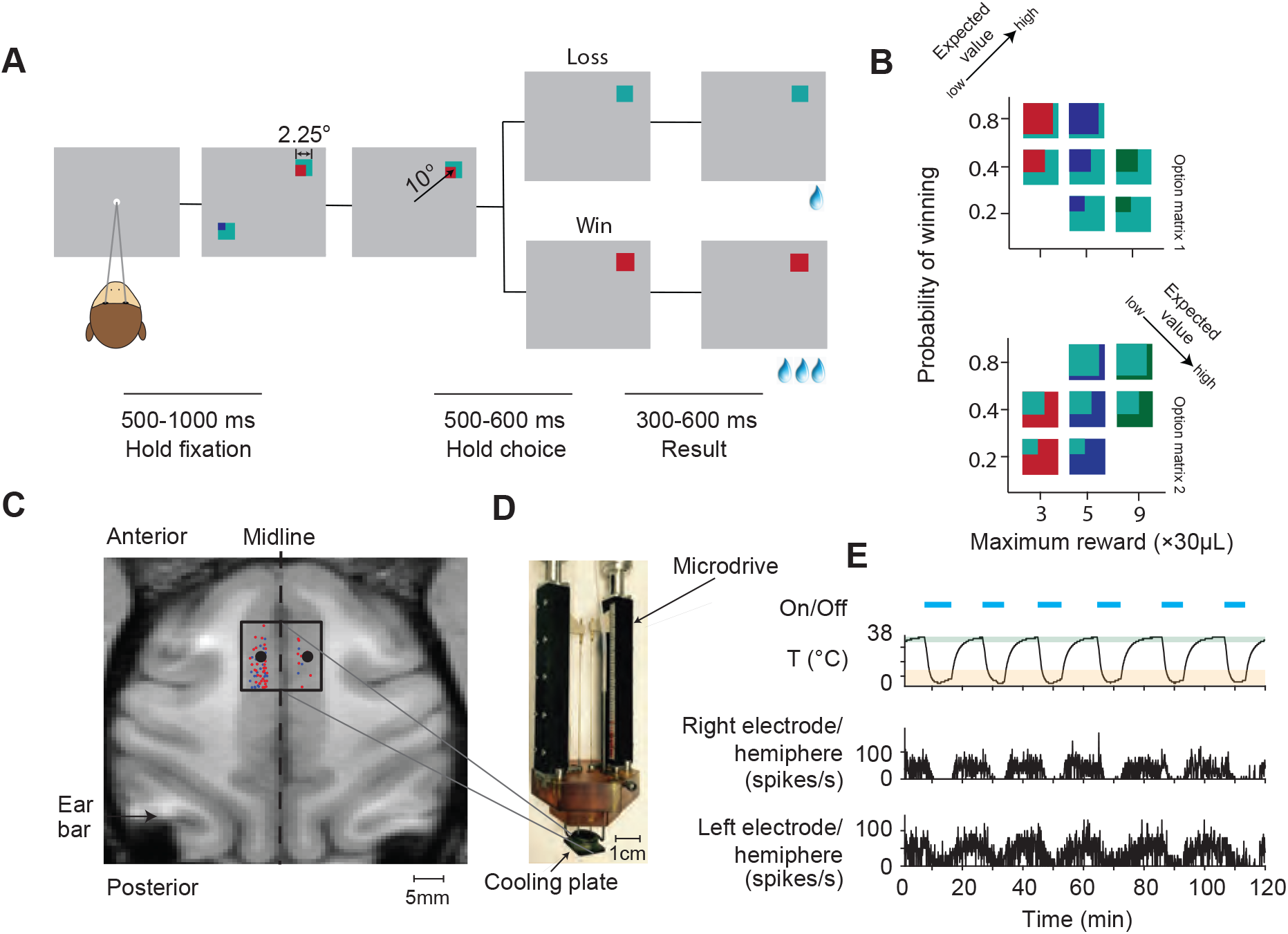
Task design and experimental setup. (A) Task design and trial sequence. Monkeys fixated a white dot while two option stimuli were presented. Monkeys could earn a reward by making a saccade to one of the option stimuli. The lines below indicate the duration of epochs in the gambling task. (B) Two sets of gamble options (option matrix 1 and option matrix 2) used in the gambling task. In a given session, we presented seven possible gamble options with three levels of maximum reward amount and three levels of winning probability (see Figure S1). Each option stimulus contains two colors. There are four different colors in total (cyan, red, blue and green) indicating four different reward amounts (increasing from 1, 3, 5 to 9 units of water, 1 unit = 30 μL of water). The proportions of the areas covered by the colors indicate the probability of receiving the corresponding reward. The expected value of the gamble targets increases along the axis indicated by the arrow (see Figure S2). (C) Cryoinactivation experiment setup. The black square in the left subplot indicates the position of the cooling plate during bilateral inactivation. The black dots within the square indicate the recording sites at which neurophysiological recordings were performed during inactivation. Red dots indicate the recording sites where task related neuronal activity was recorded in separate experiments. Blue dots indicate the recording sites with no task related neuronal activities. The back square indicates the cooling plate covers the majority of the cortical area with task related activity. (D) The cooling device consist of three parts: the cooling plate (10 mm × 12 mm), the brown plastic cap, which stabilizes the whole cooling device in the recording chamber, and the two micro-drives, which hold two tungsten electrodes monitoring the neuronal activity during inactivation. (E) A representative experimental session. The first row shows the on- and offset of the cooling device. The second row shows the temperature recorded at the cooling plate, right above the dura. The shaded green area indicates the temperature range defined as the control state, and the shaded orange area indicates the temperature range defined as the inactivation state. The third and fourth row shows the multi-unit spiking activities recorded simultaneously in both left (the third row) and right (the fourth row) SEF (see Figure S2).

The monkeys used the gamble cues in an economically rational way. They consistently selected gambles with higher reward amount (error rates: Monkey A: 9.22%; Monkey I: 1.99%; Figure 2A) and higher winning probability (error rates: Monkey A: 4.94%; Monkey I: 2.75%; Figure 2A) when the other attribute was matched. Overall, the monkeys clearly preferred options with higher expected value (EV) (Figure S1 A, B, H, and I). The monkeys were also risk-seeking, consistent with many previous studies [6,10,19,20]. For gambles with identical EV, both monkeys preferred the gamble option with the higher outcome variance, i.e. higher risk (Figure 2B, t-test, Monkey A: P(choose more risky option)=79.01%, t-test: *p*=2.37×10^-6^; Monkey I: P(choose more risky option)=73.40%, t-test: *p*=5.19×10^-4^).

**Figure 2.**
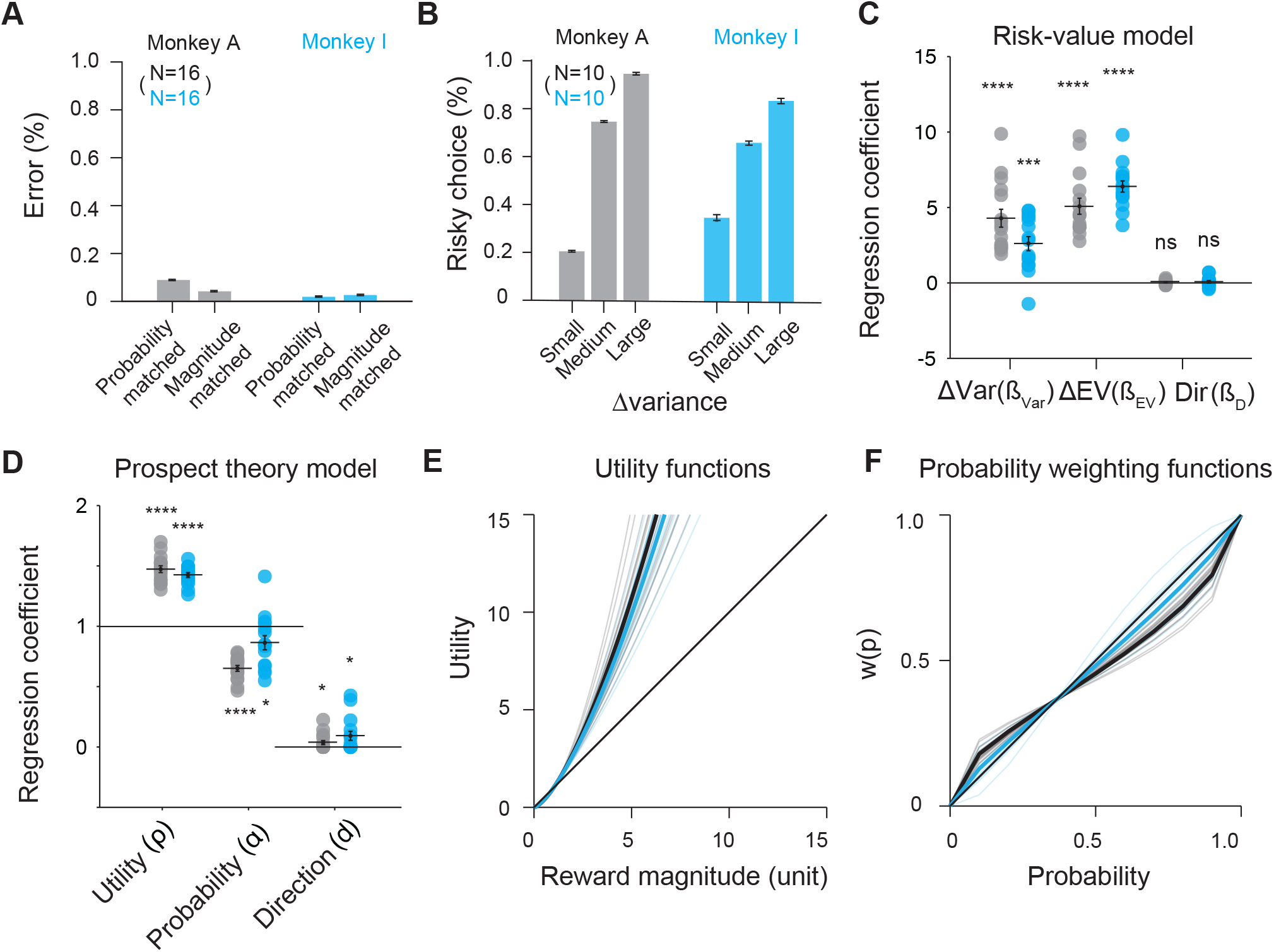
Monkeys show risk-seeking behavior. (A) The monkeys showed first-order stochastic dominance. They strongly preferred the gamble option with the higher probability or higher amount, when the other factor (reward amount or probability) was held constant. (B) The monkeys were risk-seeking. They strongly preferred the gamble option with higher variance, when expected value was identical for both gambles options. This preference increased with increasing variance differences. (C) The risk-value model explains choices between two gamble options as a function of outcome variance differences (ΔVar), expected value differences (ΔEV), and directional bias (Dir). The regression coefficients for ΔVar and ΔEV for both monkeys are significantly different from 0 (t-test, *p* <10^-4^). (Monkey A: black; Monkey I: blue). The Var coefficients are positive, indicating attraction to risk. The regression coefficients for Dir are not significant different from 0. Each dot represents an estimated regression coefficient for one experimental session. (D) The prospect theory model explains choices between two gamble options as a function of the non-linear utility of outcomes, weighted by a non-linear probability function, and the directional bias. The estimated coefficients for both utility and probability distortion are significantly different from 1 (t-test, *p* <10^-4^). The estimated coefficients for direction is slightly above 0 (t-test, *p* <0.05). (E) The estimated power utility functions for both monkeys. The thin lines denote the individual session estimations, while the thick lines denote the average estimation. The utility functions are convex, indicating risk-seeking. (Monkey A: black; Monkey I: blue). (F) The estimated probability weighting function using the 1-parameter Prelec weighting function for both monkeys. The color scheme is similar to f. Error bars denote s.e.m; *\*,p* <0.05; *\*\*\*\*,p* <10^-4^ See also Figure S1 for choice patterns with both option matrices separately.

We quantified the monkeys’ risk preference using two standard economic models: the risk-value and the prospect theory model (Methods). The risk-value model is derived from financial theory and decomposes the subjective value of each option into a weighted linear combination of EV and variance risk, computed as the variance (Var) of the gamble outcomes [10,16,17]. It outperformed models using only the EV or the Var term, and models using coefficient of variance of the gamble outcomes, an alternative measure of risk [22] (Table S2). The monkeys preferred options with higher EV (Figure 2C, t-test, Monkey A: 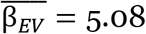, *p*= 5.60×10^-8^; Monkey I: 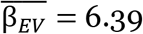, *p*= 8.03×10^-11^) and higher Var, leading to risk seeking behavior, (Figure 2C, t-test, Monkey A: 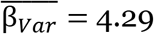, *p*= 2.03×10^-6^; Monkey I: 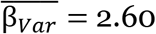, *p*= 7.00×10^-5^).

The prospect theory model is derived from expected utility theory and estimates subjective value using a non-linear utility and probability weighting function [9,22,23]. This model predicted the monkeys’ choice behavior better than models using either utility or probability weighting functions alone, and also slightly better than the risk-value model (Table S2). The best-fitting utility functions of both monkeys were convex (Figure 2 D and E, t-test, Monkey A: 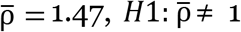, *p*= 2.39×10^-11^; Monkey I: 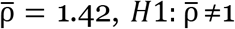, *p*= 6.30×10^-12^). In addition, the monkeys also significantly overweighed low and underweighted high probabilities (Figure 2 E and F, t-test, Monkey A: 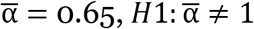, *p*= 2.32×10^-10^; Monkey I: 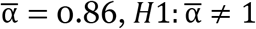, *p*= 0.03). Therefore, the monkeys were attracted disproportionally to large reward amounts and overestimated the likelihood of obtaining them when the winning probability was low, leading to risk seeking behavior.

Thus, both economic models indicated a strong preference for riskier options. In contrast, there was only very weak evidence for directional bias (risk-value model: t-test, Combined: 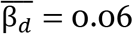, *p*=0.21; Monkey A: 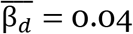, *p*=0.23; Monkey I: 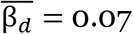, *p*=0.42; prospect theory model: t-test, Combined: 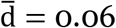, *p*=0.01; Monkey A: 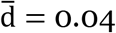, *p*=0.05; Monkey I: 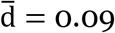, *p*=0.03). There was no evidence that the monkeys tended to repeat the previous choice direction (t-test, Combined: 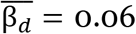, *p*=0.21; Monkey A: 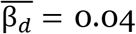, *p*=0.23; Monkey I: 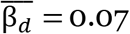, *p*=0.42).

### SEF inactivation reduces risk-seeking

SEF neurons encode action value signals that reflect the subjective value of options in the oculomotor gambling task and are correlated with the monkeys’ choices [8]. To test if these signals have a causal effect on decision making, we examined whether bilateral inactivation of SEF influenced monkeys’ behavior in the oculomotor gambling task, using a cryoplate (Figure 1 C and D). This method allows us to quickly and reversibly inactivate the SEF in both hemispheres [24]. We monitored neuronal activity in both SEF hemispheres during control and inactivation conditions. Consistent with previous reports [25,26], the spiking activity decreased with decreasing temperature in both hemispheres (Figure 1E and Figure S2). Neuronal activity was less affected as distance increased between the recording sites and the cooling plate (Figure S2), so that the cooling effect was restricted to SEF. In total, we performed 31 bilateral inactivation sessions (Monkey A: 16 sessions and Monkey I: 15 sessions), with an average of 1399 successful trials and 7 periods of inactivation per session.

The effect of SEF inactivation on risky choice was highly consistent across the two monkeys. During inactivation, we observed in both monkeys some small changes in saccade metrics (Figure S3), fixation stability (Figure S3) and reaction times (S4A and Table S3), consistent with previous findings [27–30]. These changes in oculomotor behavior were too small to affect choice. SEF inactivation caused only small and inconsistent changes in error rate when the options only differed in either winning probability or magnitude (Figure 3A). Therefore, SEF inactivation did not affect the ability of the monkeys to use the visual cues for economically rational choices. Nevertheless, both monkeys showed a significantly altered pattern of choice during SEF inactivation: they were consistently less risk seeking (Figure 3 C and E). The monkeys showed reduced risk preference in 90% (28/31) of inactivation sessions as measured by the risk-value and prospect theory models.

**Figure 3.**
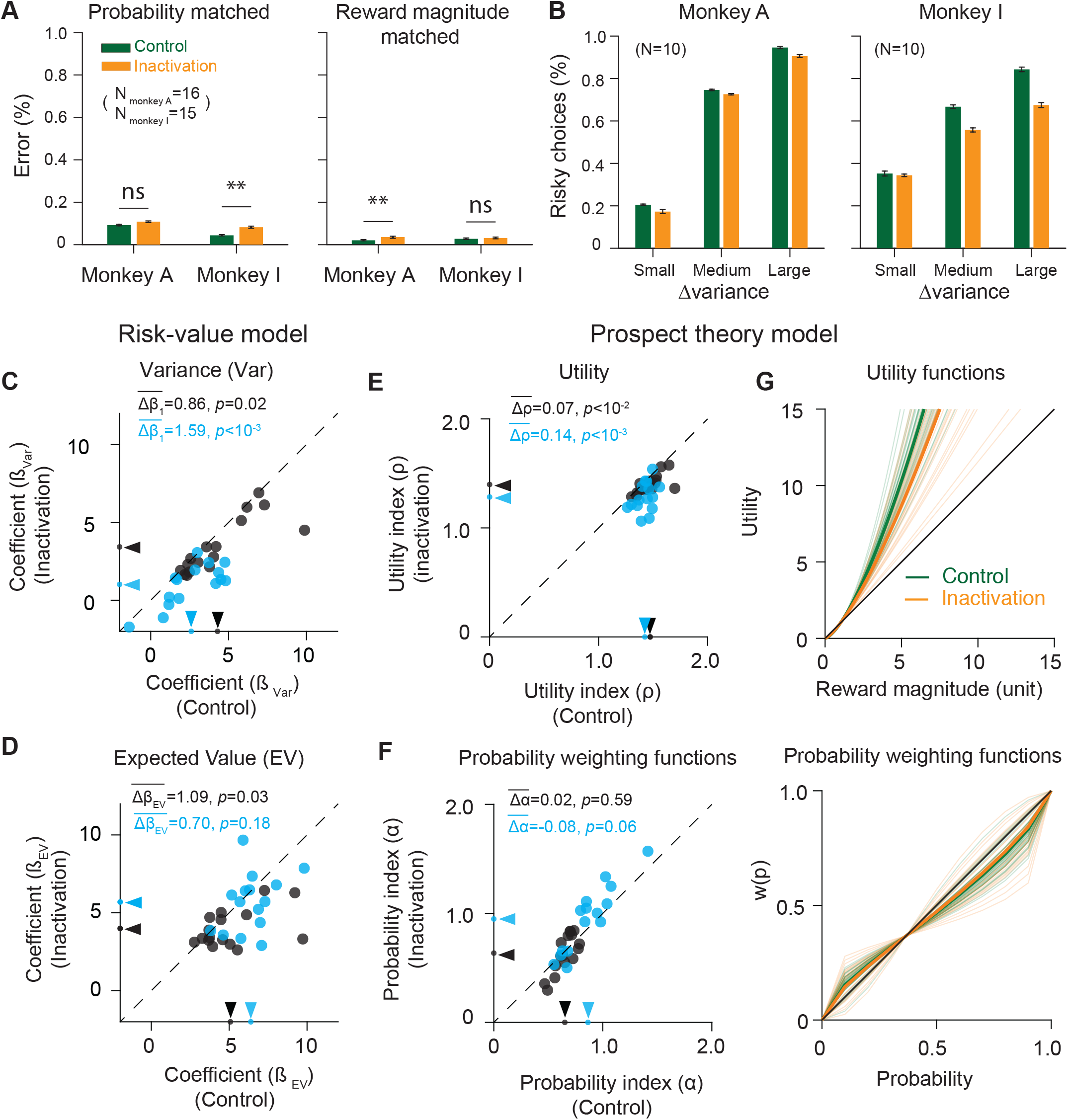
SEF inactivation reduces risk-seeking. (A) Inactivation has only a small and non-consistent effect on the monkeys’ ability to choose the optimal gamble option, if they vary only in one factor (either magnitude or probability). Error rates are low both in the control (green) and inactivation (orange) condition and do not show consistent differences across monkeys. (B) Inactivation reduces risk-seeking. The preference for gamble options with larger outcome variance is less pronounced during inactivation (orange) compared to the control (green) condition (paired t-test, Combined: *p*=1.46×10^-3^; Monkey A: *p*=0.05; Monkey I: *p*=0.01). (C-D) Reduction of risk-seeking estimated by the risk-value model. (C) Comparison of the coefficients for risk (Var) in the control and inactivation condition for all experimental sessions (Monkey A: black; Monkey I: blue). The arrows indicate the mean value for each condition. The coefficients were consistently lower in the inactivation condition, indicating reduced preference for risk. (D) The coefficients for expected value (EV) were slightly, but consistently decreased in the inactivation condition. All conventions are identical to C here and in all other scatter plots. (E-G) Reduction of risk-seeking estimated by the prospect theory model. (E) The parameter controlling curvature of the utility function is consistently decreased in the inactivation condition, which indicates reduced risk-seeking. (F) The parameter controlling probability weighting is not significantly changed during inactivation. (G) The corresponding utility functions (top) and probability weighing functions (bottom) are shown for both monkeys during the control (green) and inactivation (orange) condition. Thin lines indicate individual sessions, while thick lines indicate the average function. Error bars denote s.e.m.; paired t-test, ns, nonsignificant; *, *p* <0.05; **, *p* <10^-2^ ***, *p* <10^-3^; ****, *p* <10^-4^. See SEF inactivation effect on saccade reaction times in Table S3. See reduction of risk-seeking estimated by the prospect theory model separately for both monkeys in Figure S4C.

In the risk-value model, the risk term (Var) coefficients were significantly smaller during inactivation compared to the control condition (Figure 3C, paired t-test, Combined: 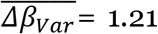, *p*= 9.92×10^-6^; Monkey A: 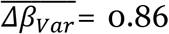, *p*= 0.02; Monkey I: 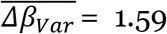, *p*= 1.23×10^-4^). Thus, both monkeys showed a strong reduction of risk preference during inactivation (Combined: 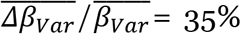; Monkey A: 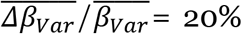; Monkey I: 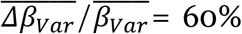). For gambles with identical EV, both monkeys chose the higher risk option significantly less often (Figure 3B, paired t-test, Combined: ΔP(choose higher risk option)= 6.97%, *p*= 1.46×10^-3^; Monkey A: ΔP(choose higher risk option)= 4.29%, *p*= 0.05; Monkey I: ΔP(choose higher risk option)= 9.66%, *p*= 0.01). The monkeys’ choices were also less determined by EV differences during inactivation (Figure 3D, paired t-test, Combined: 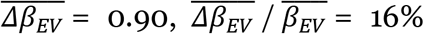, *p*= 0.01; Monkey A: 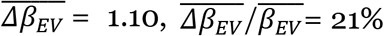, *p*= 0.01; Monkey I: 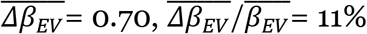, *p*= 0.18). Across all trials, the monkeys chose the smaller EV option significantly more often (paired t-test, Combined: ΔP(choose lower EV option)= 1.71%, *p*= 3.63×10^-4^; Monkey A: ΔP(choose lower EV option)= 1.46%, *p*= 5.39×10^-3^; Monkey I: ΔP(choose lower EV option)= 1.97%, *p*= 0.02). However, this effect was less pronounced than the one resulting from the lower preference for risk (Figure S4B).

In the prospect theory model, the utility functions of both monkeys were less convex during inactivation (Figure 3 E and G, paired t-test, Combined: 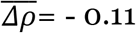, *p*=6.32×10^-6^; Monkey A: 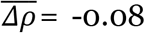, *p*= 2.00×10^-3^; Monkey I: 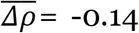, *p*=7.35×10^-4^). The monkeys showed less overestimation of high reward amounts during inactivation. In contrast, the probability weighting function, which captures the monkeys’ estimation of the probability of winning, remained unchanged (Figure 3 F and G, paired t-test, Combined: 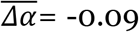, *p*= 0.17; Monkey A: 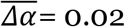, *p*= 0.59; Monkey I: 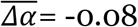, *p*= 0.06).

We tested if changes in motor strategies could explain this preference change, since manipulation of dopaminergic receptors in frontal eye field can change positional bias and the tendency to repeat actions [31]. There was no significant change of directional preference (risk-value model: t-test, Combined: p=0.19; Monkey A: p=0.06; Monkey I: p=0.01; prospect theory model: t-test, Combined: p=0.31; Monkey A: p=0.06; Monkey I: p=0.23) or repetition of the previous choice direction (Combined: p=0.21; Monkey A: p=0.91; Monkey I: p=0.16). The reduced risk-seeking reflects therefore a true change in choice preference.

### SEF inactivation increases trial desertion after gamble loss

During the result epoch, SEF neurons encode reward prediction error (RPE), the difference between expected and actual reward [6]. RPE signals are thought to guide reinforcement learning and updating of action value signals [6,31]. We therefore tested whether SEF inactivation influenced the monkeys’ sensitivity to these locally encoded RPE signals. Following the loss of a gamble, both monkeys occasionally actively broke fixation by making a saccade outside of the fixation window (Figure S3C), thus deserting the trial before reward delivery. This behavior was maladaptive, because it did not change the outcome of the trial, and substantially prolonged the time until reward delivery, as well as the time until the next chance to make a choice. Trial desertion developed spontaneously, was sensitive to negative RPE, and increased with larger errors (Figure 4A). Interestingly, desertion rate was significantly higher in choice trials than no-choice trials (paired t-test, Combined: 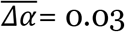, *p*= 4.73×10^-4^; Monkey A: 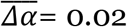, *p*= 0.02; Monkey I: 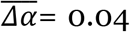, *p*= 0.01). Following SEF inactivation, both monkeys were substantially more sensitive to RPE in choice trials (Figure 4 A and B top, paired t-test, Combined: 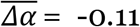, *p*= 9.35×10^-9^; Monkey A: 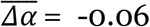 *p*= 1.82×10^-4^; Monkey I: 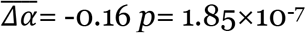), but not in no-choice trials (Figure 4 A and B bottom, paired t-test, Combined: 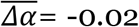, *p*= 0.06; Monkey A: 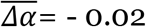, *p*= 0.16; Monkey I: 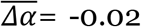, *p*= 0.21.) In addition, desertion rates also significantly increased in all other task epochs of choice trials during inactivation (Figure S4). Thus, outcome monitoring signals in the SEF were not necessary to drive desertion behavior. On the contrary, SEF activity seems to be necessary to suppress desertion behavior throughout the task, but in particular following aversive events following free choices (Figure 4 A and B).

**Figure 4.**
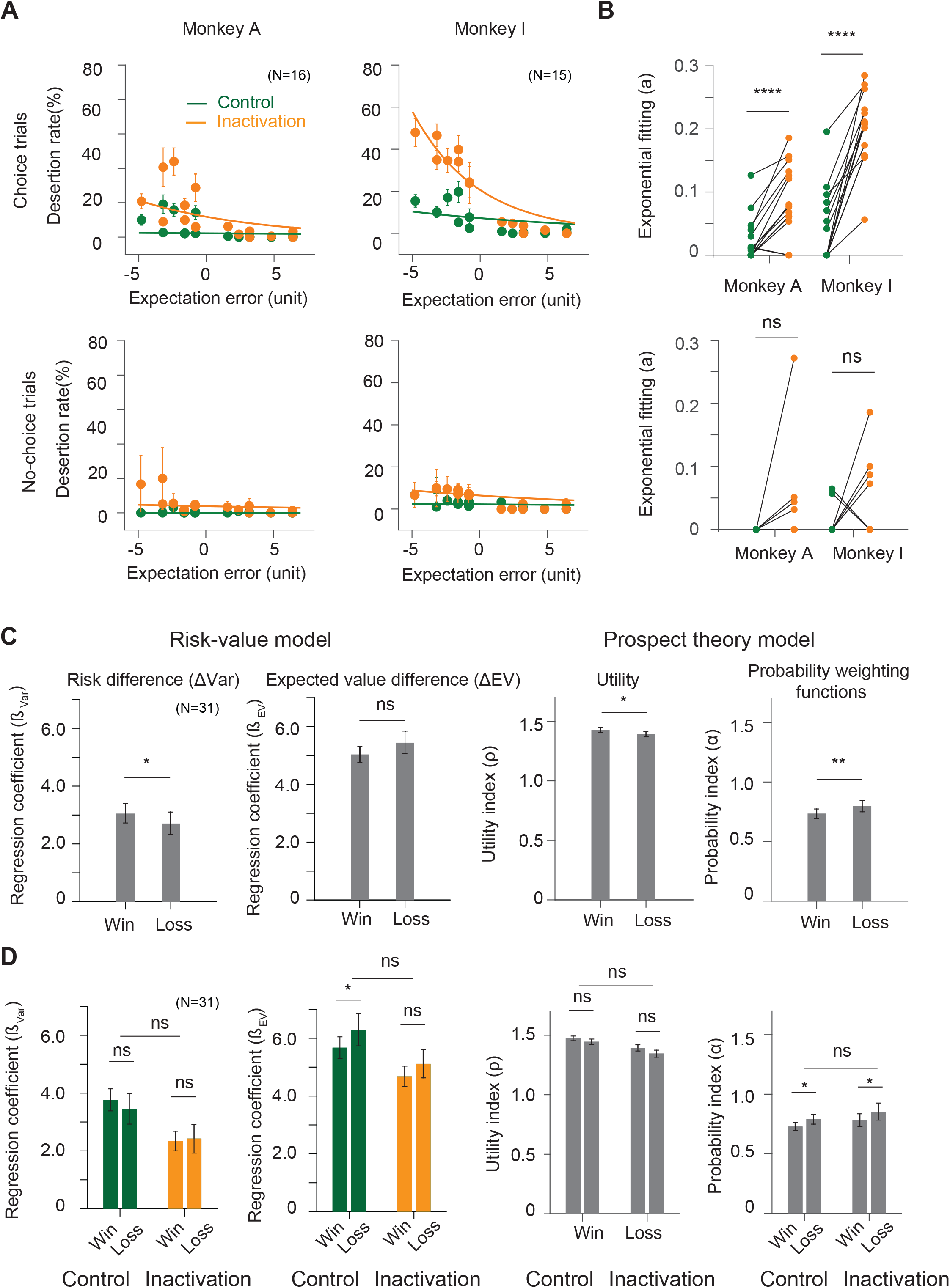
Influence of SEF inactivation on desertion rates and gamble history effect. (A) Desertion rates during result periods as a function of reward expectation errors, the differences between actual and anticipated reward, during control (green) and inactivation (orange) condition in choice trials (top) and no-choice trials (bottom). The overall desertion rates are estimated using an exponential fit, indicated by the colored lines. These rates are significantly increased during inactivation in choice trials. This increase is larger with larger negative prediction errors. Trial desertion here is defined as making saccades actively outside of the fixation windows (see Figure S3C and D). See Figure S4 for trial quitting rates in other task epochs. (B) Changes of scaling parameters of the exponential functions for each individual experiment session in choice trials (top) and no-choice trials (bottom). Each line shows the change in the scaling parameters in control and inactivation condition from one experiment session. (C) Changes of risk preference based on gamble outcome history across trials during both control and inactivation conditions. The monkeys were less risk-seeking following a lost gamble as compared to a won gamble. This effect is captured both by the risk-value (left two plots) and the prospect theory (right two plots) models. In the risk-value model, the risk coefficient is significantly lower on trials following a gamble loss, while the expected value coefficient stays the same. In the prospect theory model, the utility function becomes less concave (*ρ* closer to 1) and the probability weighting function becomes more linear (*α* closer to 1) on trials following a gamble loss. (D) The effect on risk preference by gamble outcome history does not change across control (green) and inactivation (orange) conditions. Error bars denote s.e.m.; paired t-test, ns, non-significant; *, *p* <0.05; **, *p* <10^-2^ ***, *p* <10^-3^, ****, *p* <10^-4^.

### SEF inactivation does not affect reward-history dependent adjustments of risk preference

Both monkeys showed a significant change of risk preference depending on the preceding gamble outcome. They were less risk seeking when they had lost the previous gamble than when they had won it. In the risk-value model, this manifested itself in a significant difference in the Var coefficient (Figure 4C, paired t-test, Combined: 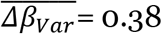, *p*= 0.03; Monkey A: 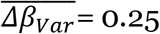, *p*= 0.10; Monkey I: 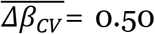, *p*= 0.09), while the EV coefficient was not significantly different (paired t-test, Combined: 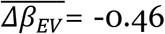, *p*= 0.11; Monkey A: 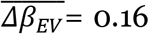, *p*= 0.45; Monkey I: 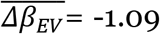, *p*= 0.14). In the prospect theory model, the same change in risk preference manifested itself in less convex utility functions (paired t-test, Combined: 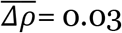, *p*= 0.05; Monkey A: 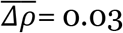, *p*= 0.06; Monkey I: 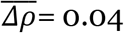, *p*= 0.27) and in a more linear probability weighting function (paired t-test, Combined: 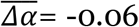, *p*= 0.01; Monkey A: 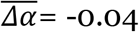, *p*= 0.09; Monkey I: 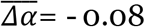, *p*= 0.01) after losing in the previous trial. Thus, following a loss, both monkeys were more risk averse in their subsequent choice. However, this gamble outcome effect persisted during SEF inactivation and did not show any significant changes (Figure 4D, paired t-test, 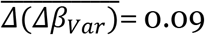, *p*= 0.77; 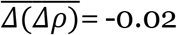, *p*= 0.62; 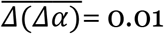, *p*= 0.82). Therefore, although gamble outcome history modulates the monkeys’ gamble value estimation, this adjustment does not depend on local RPE signals in SEF.

## Discussion

Decision related activity has been observed in many brain regions [32]. However, it remains unknown whether this activity is causally related to the decision process [33–35]. Here we showed that the SEF, an oculomotor area within the medial frontal cortex, does play a causal role in regulating risky and impulsive behavior in oculomotor decisions.

SEF is only one among a number of cortical [1,13,15,36] and subcortical [14,37–39] brain areas that contribute to decision-making under risk. However, the effect of SEF inactivation is not a simple decrease in decision accuracy, as would be expected if SEF operates in parallel with other areas that contain redundant signals, so that SEF inactivation merely reduces the overall strength of the decision variable. Instead, SEF seems to selectively mediate the effect of risk preferences, but not expected value, on choice. Eliminating these signals cannot be fully compensated for by other parts of the decision-making circuit.

Risk preference is often seen as a fundamental, stable personality trait [40]. However, the risk preference of individuals can vary substantially across different behavioral domains [41]. Even when tested only within the financial domain, risk preference varies [42]. These findings suggest that risk preference is not a stable personality trait, but rather emerges during decision-making in a context-dependent manner. Risk-attitude depends on beliefs about the environment, the set of available options, and the contingencies governing action outcomes [43]. In the context of our experimental task, the small stakes and large number of trials likely reduced the averseness of losing a gamble and thus induced risk-seeking behavior [42,43]. These contextual factors are not directly observable and must be inferred. Nevertheless, they are important elements of a cognitive representation of task space [46]. A recent study [43] shows that rhesus monkeys show very different utility and probability weighting functions when tested with different gamble tasks. This supports the hypothesis of the use of flexible cognitive processes in constructing risk attitudes in a context-dependent fashion.

A number of cortical areas might be important in influencing risk-attitude. Orbital frontal cortex (OFC) is involved in representing task space [47,48] and contains risk selective neurons [10,36]. Recent lesion experiments in macaques indicate also a role of ventrolateral prefrontal cortex (VLPFC) in learning and encoding the probability of reward outcomes [49]. In addition, ACC has also been shown to be correlated with risk uncertainty [12–14]. SEF receives synaptic input from frontal areas including OFC, VLPFC, and ACC, and projects to the frontal eye field, and superior colliculus [50]. It integrates sensory and task context information to guide the selection of appropriate actions [5]. Thus, the effect of SEF inactivation likely reflects the diminished influence of these belief states about the task structure, so that the subjective value of a gamble option is less determined by risk preference.

Perturbations of dopaminergic activity can also modulate risky choices [37,51–53]. In rodents, ventral tegmental area stimulation after non-rewarded choices increased subsequent willingness to choose a risky gamble [51]. In contrast to modulating risk preference by changing cognitive processes, these perturbations likely change choice behavior by modulating EV updates using model-free learning mechanisms. The fact that SEF inactivation does not affect reward-history dependent EV adjustments suggests the independent contributions of two different brain circuits to the evaluation of uncertain reward options: risk preference is associated with a goal-directed frontal cortex-based circuit, including SEF, while EV representation is associated with a more automatic subcortical circuit.

The monkeys sometimes desert the trial following an unexpected loss. This behavior likely represents an automatic response to the aversive outcome, especially following free choices. The fact that SEF inactivation increased this behavior, but only during free choice trials, cannot be explained by the fixation quality during inactivation (Figure S3 and Methods). Instead, it suggests that SEF activity contributes to self-control by suppressing automatic, but maladaptive, responses and promoting behavior that maximizes long-term reward. Such a role would be consistent with the well-known contribution of SEF to other forms of executive control [5,54].

In conclusion, our results demonstrate for the first time the causal role of SEF in mediating the effect of risk preference on decisions under uncertainty. These findings provide new insight into the neuronal circuits underlying inconsistent, context dependent choices under risk observed across humans [18,42,55] and non-human primates [45], and may provide the basis for more effective treatments of highly maladaptive impulsive risky behaviors.

## Acknowledgements

The authors would like to thank Xiaoqin Wang, Marcus Jeschke, Bill Nash and Bill Quinlan for technical help, Tirin Moore and William Newsome for invaluable scientific discussion, and Christof Fetsch, Marc Zirnsak, Donatas Jonikaitis, and Megan Wang for comments on the manuscript. This work was supported by the National Institutes of Health through grants 2R01NS086104 and 1R01DA040990 to VS.

## Author contributions

X.C. and V.S. formulated the hypotheses, designed the experiment, analyzed the data, and drafted the manuscript; V.S. secured funding; X.C. carried out the experiment and collected the data.

## STAR Methods

### KEY RESOURCES TABLE

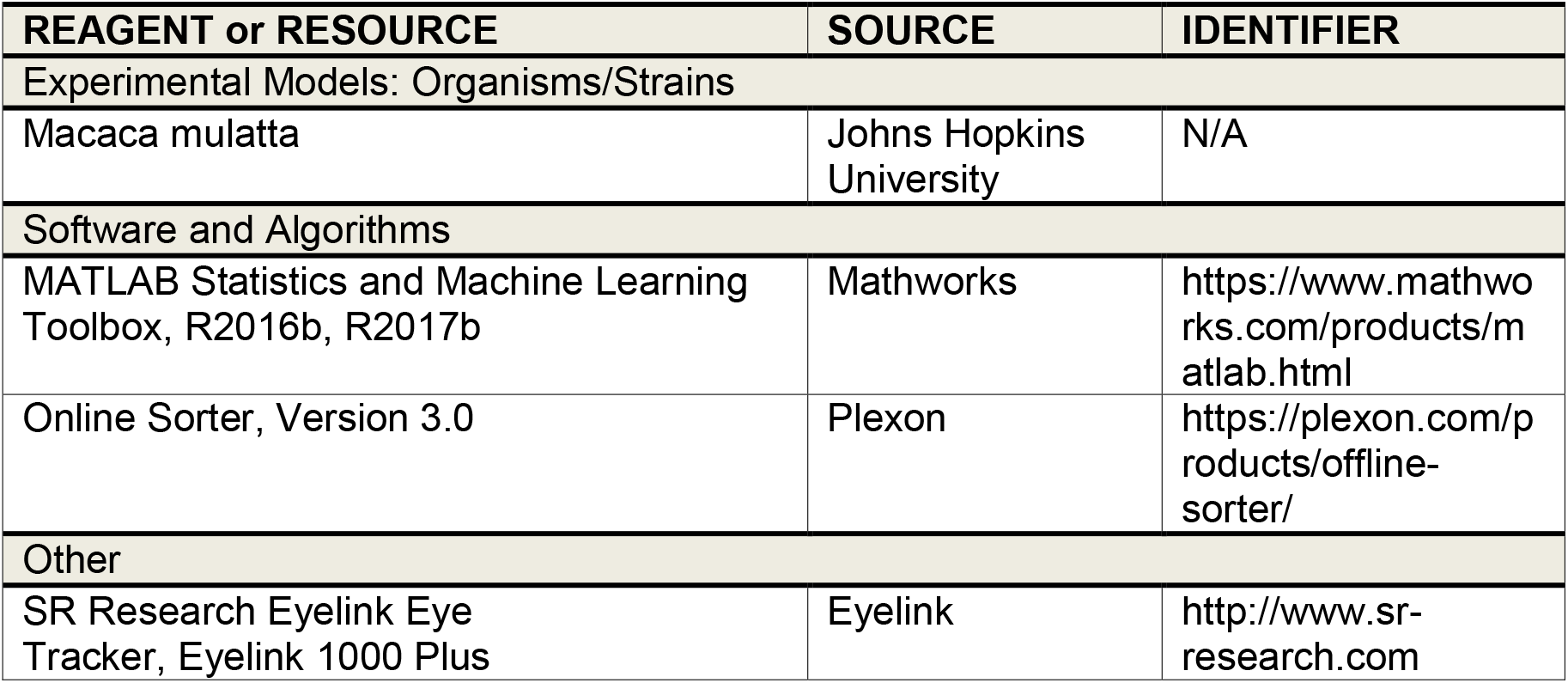

### CONTACT FOR REAGENT AND RESOURCE SHARING

Further information and requests for resources should be directed to and will be fulfilled by the Lead Contact, Veit Stuphorn.

### EXPERIMENTAL MODEL AND SUBJECT DETAILS

All animal care and experimental procedures were in compliance with the US Public Health Service policy on the humane care and use of laboratory animals, and were approved by Johns Hopkins University Animal Care and Use Committee. Two male rhesus monkeys (Macaca mulatta, Monkey A: 7.5 kg, Monkey I: 7.2 kg) were trained to perform the tasks used in this study. After training, we placed a hexagonal chamber (29 mm in diameter) centered over the midline, 28 mm (Monkey A) and 27 mm (Monkey I) anterior of the interaural line.

### METHOD DETAILS

#### Electrophysiological techniques

During each bilateral inactivation session, single units were recorded using two tungsten microelectrodes with an impedance of 2-4 MΩs (Frederick Haer, Bowdoinham, ME), one in each hemisphere (Figure 1C). The microelectrodes were advanced, using a self-built microdrive system. Data were collected using the PLEXON system (Plexon, Inc., Dallas, TX). The electrodes penetrated the cortex perpendicular to the surface of the SEF. The depths of the neurons were estimated by their recording locations relative to the surface of the cortex.

#### Behavioral task

In the task, the monkeys had to make saccades to peripheral targets that were associated with different reward amounts and probabilities (Figure 1A). The targets were colored squares, 2.25×2.25^0^ in size. They were always presented 10° away from the central fixation point at a 45, 135, 225, or 315^0^ angle. There were 7 different gamble targets (Figure 1B), each consisting of two colors corresponding to the two possible reward amounts. The portion the target filled with each color corresponded to the probability of receiving the corresponding reward amount. Four different colors indicated four different reward amounts (increasing from 1, 3, 5 to 9 units of water, where 1 unit equaled 30 μl). The minimum reward amount for the gamble option was always 1 unit of water (indicated by cyan), while the maximum reward amount ranged from 3 (red), 5 (blue) to 9 units (green), with three different probabilities of receiving the maximum reward outcome (20, 40, and 80%). Only gamble options from either option matrix1 or option matrix 2 were used in an experimental session.

The task consisted of two types of trials - choice and no-choice trials. All trials started with the appearance of a fixation point at the center of the screen (Figure 1C), on which the monkeys were required to fixate for 500-1000 ms. In choice trials, two targets appeared in two locations that were randomly chosen from across the four quadrants (resulting in 12 distinct possible spatial configurations for each pair of gamble options). Simultaneously, the fixation point disappeared, which indicated to the monkeys that they were now free to choose between the gambles by making a saccade toward one of the targets. Following the choice, the non-chosen target disappeared from the screen. The monkeys were required to keep fixating on the chosen target for 500-600ms, after which the gamble outcome was revealed. The two-colored square changed into a single-colored square associated with the final reward amount. The monkeys were required to continue to fixate on the target for another 300 to 600 ms, during which the result cue was still displayed, until the reward was delivered. We observed quantitatively similar results using both option matrices (Figure S1). We therefore report the combined results in the manuscript. All 7 gamble options in each option matrix were systematically paired with all other options from that matrix. This resulted in 21 different combinations of gamble options in choice trials. The sequence of events in no-choice trials was the same as in choice trials, except that only one target was presented. In these trials, the monkeys had to make a saccade to the given target in order to get the fluid reward.

If the monkey deserted a choice trial before choosing between the gambles, the choice trial was simply repeated. However, if the monkey deserted the trial after the choice, but before reward was delivered, the next trial was an unscheduled no-choice trial. The target shown on this no-choice trial depended on the stage at which the monkey had deserted the preceding trial. If the monkey had deserted before the gamble result was revealed, the target was the previously chosen gamble option. If the monkey had deserted the trial after the gamble result was shown, the target was the previously indicated sure reward that was the gamble outcome. Thus, a gamble option or result was binding, once it was chosen or revealed, respectively. Accordingly, desertion behavior was suboptimal and only reduced the average reward rate across trials.

#### Cryogenic inactivation apparatus and procedure

To determine the location of the SEF, we obtained magnetic resonance images (MRI) for Monkey A and Monkey I. We used the location of the branch of the arcuate sulcus as an anatomical landmark. Before the inactivation experiment, we identified the SEF by neurophysiology recordings (Figure 1C). In both monkeys, we found neurons active during the saccade preparation period in the region from 0 to 11 mm anterior to the genu of the arcuate branch and within 5 mm to 2 mm of the longitudinal fissure. We designated the cortical areas with saccade-preparation related activity as belonging to the SEF [8], consistent with previous studies from our lab and existing literature [44,56].

Cooling plates (Figure 1C) were used to inactivate the SEF bilaterally (10mm from anterior to posterior and 12mm from left to right). This method allows us to rapidly and repeatedly inactivate a large and confined surface cortical area [24,57]. The cooling method followed the design by Lomber et al. [24]. Room temperature methanol was pumped through Teflon tubing that passed through a dry ice bath, in which it was reduced to subzero temperature. The chilled methanol was then pumped through a cryoloop attached to a stainless-steel plate placed over the dura, which cooled down the underlying cortical tissue. The methanol was then returned to the same reservoir from which it came to form a closed loop. The cortical temperature on the dura was monitored by a micro-thermocouple attached to the cooling plate. At the same time, two electrodes recorded cortical activity in the left and right hemisphere. During each session, monkeys initially performed the task for 10-15 min in the control state. Then the SEF region was deactivated bilaterally for 10-15 min by pumping chilled methanol through the cryoloop while the task continued. The cortical temperature returned quickly to normal after switching off the methanol pump (Figure 1E), while the monkey continuously performed the task. This whole process was repeated throughout the experimental session and resulted on average in 1399 successful trials, which is on average 7 repetitions of control/inactivation cycles. In the control state, the temperature measured at the cooling probe was 35-39 °C. During the inactivation state, the temperature at the cooling plate was reduced to 0-15 °C. Transition trials, right after turning on the pump and turning off the pump, with the temperature between 34 and 16 °C, were not used in the behavioral analysis. The monkeys were sitting in an acoustic noise-isolated chamber. The methanol pump was placed outside this chamber.

### QUANTIFICATION AND STATISTICAL ANALYSIS

In general, two-tailed t-tests were used for statistical tests, unless specified otherwise.

#### Risk behavior analysis

Trial-by-trial data was collected during control and inactivation. We quantified the monkeys’ risk behavior using two types of risk models: risk-value models and prospective theory models. All reported *p* values regarding mean differences between control and inactivation conditions are results of two-tailed paired t-tests. P values relating to gamble history effects are based on one-tailed paired t-tests.

The **risk-value model** is derived from financial theory [16] and represents the value of a gamble as the sum of multiple terms related to the distribution of possible gamble outcomes. The first term is the mean value of the gamble outcome distribution (i.e., the expected value of the gamble). The second term is the variance of the gamble outcomes (i.e., variance risk). The sign of this term determines if outcome variance increases (risk-seeking) or decreases (risk-averse) the value of gambles. In the following, we will refer to this second component simply as risk. In more complex models of this type, higher statistical moments describing the outcome distribution (skewness, kurtosis) are also taken into account. However, here we will not use these higher-order terms.

We used logistic regression to quantify the ability of the risk-value model to predict choice behavior. We assumed that choice depended in a stochastic fashion on the difference in subjective value between the two gamble options. We used a soft-max decision function to model this aspect of behavior:

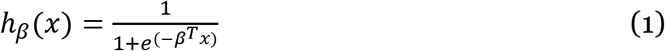

such that:

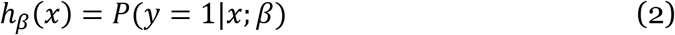

where y ∊ {0,1} is a dummy variable indicating whether the monkeys choose the first option or not, and *β* is the set of weights learned by the model. The first option is defined as the left option if the choice options were on both left and right visual field, and is defined as the up option if the choice options were both on the same visual field. The full risk-value model has two terms: expected value (EV) and outcome variance (Risk, Var). Expected value is defined as the arithmetic mean of the outcomes: EV = V_*win*_ × *p_win_* + V_*loss*_ × *p_loss_*, with *V_win_* denoting the winning reward magnitude, *V_loss_* denoting the losing reward magnitude, *p_win_* denoting the winning probability, and *p_loss_* denoting the losing probability. We defined risk as the variance of the gamble option [17]: 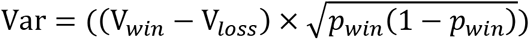 and coefficient of variance [21]: 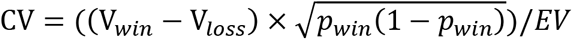 (Table S1). We achieved slightly better behavioral fitting by using standard deviation than coefficient of variance (Table S2). We tested three variants of the risk-value model, whereby subjective value of a gamble depended only on: (3) expected value, (4) risk, (5) or both (the full model):

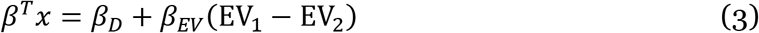

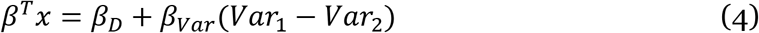

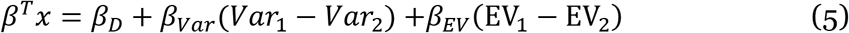

Comparing the predictions of the three versions with trial-by-trial choices of the monkey allowed us to determine, if both factors were necessary to predict choice behavior. In order to quantify the tendency of choosing the same direction as in the previous trial, we added an additional parameter which represent whether the choice option appear at the previous chosen direction or not for both choice options.

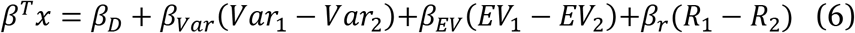

*R_i_* is 1 if the location of the option i is the same as the chosen direction in the previous trial, and *R_i_* is 0 if the location of the option i is different.

We used the gradient descent algorithm to minimize the cost function, which represents negative log-likelihood function, over training examples:

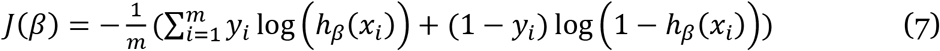

Both EV and Var were normalized to [0, 1] to enable the comparison among different independent variables. *β_D_* represents the directional bias. The regression coefficient for Var indicates the risk attitude. A negative sign of the coefficient indicated that increased outcome variance reduced subjective value, indicating risk-aversion, while a positive sign indicated risk-seeking.

**Prospect theory** is derived from classical expected value theory in economics [18] and assumes that the subjective value of a gamble depends on the utility of the reward amount that can be earned, weighted by the ‘subjective’ estimation of the probability of the particular outcome. Both the utility function and the probability function can be non-linear and thus might influence risk preference. Prospect theory also makes the assumption that utilities are perceived in a relative framework (i.e., as gains or losses relative to a reference point), not an absolute framework (i.e., the total amount of earned reward). However, this aspect of the model is irrelevant for our study, because the monkey does not encounter negative outcomes, so that for each individual trial the relative and absolute reference frame make identical predictions.

We assumed again a soft-max decision function where the probability of selecting the gamble was indicated by the difference of the subjective value of the two options:

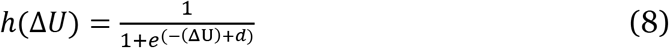

where Δ*U* = *U*_1_ – *U*_2_ is the utilities difference between gamble options, and *d* is the directional bias between two options. The utility of the choice option *i* was calculated as following:

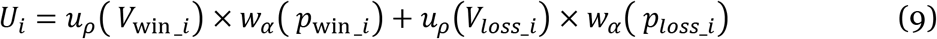

where *u_ρ_*(*V*) is a power function to model the utility function, following previous research [9,22]:

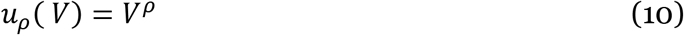

and *w_α_*(*p*) is a 1-parameter Prelec function to model the probability weighting function, as commonly done [9,22,23,58]:

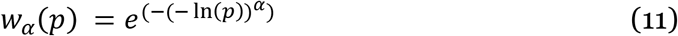

*ρ* in equation (10), *α* in equation (10) and *d* in equation (8) were free parameters optimized by a Nelder-Mead search algorithm to minimize the sum of negative log likelihoods with respect to the utility function. As in classical expected value theory in economics [22], a convex utility function (*ρ* > 1) implies risk seeking, because in this scenario, the subject values large reward amounts disproportionally more than small reward amounts. Gain from winning the gamble thus has a stronger influence on choice than loss from losing the gamble. In the same way, a concave utility function (*ρ* < 1) implies risk seeking, because large reward amounts are valued disproportionally less than small ones. Independently, a non-linear weighting of probabilities can also influence risk attitude. For example, a S-shaped probability weighting function (α < 1) implies that the subject overweighs small probabilities and underweights the large probabilities. This would lead to higher willingness to accept a risky gamble, because small probabilities to win large amounts would be overweighted relative to high probabilities to win moderate amounts.

As with the variability risk model, we tested three variants of the prospect theory model: 1) a full model, in which both utility function and weighting function were allowed to be non-linear, 2) a ‘utility-only’ version, in which only the utility function was allowed to be non-linear, and 3) a ‘probability weighting only’ version, in which only the probability weighting function was allowed to be non-linear.

#### Model comparison

The Bayesian information criterion [59,60] was used for model comparison.

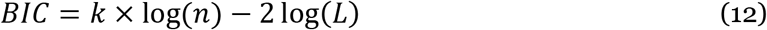

where log(*L*) is the log-likelihood (*LL*) of the model, n is the number of trials. k is the number of free parameters to be estimated.

In addition, we also combined all the trials across different experiment sessions from one monkey in a given task together. We then performed five-fold cross-validation method with different models based [43]. During cross-validation, we randomly divided all the trials into training set (80%) and test set (20%). We used training set to optimize the parameters for a given model, and use the test set to calculate LL to evaluate the model (Table S3). Cross validation procedures were repeated 50 times independently for each monkey per task.

#### Desertion Behavior

We used an exponential function to quantify the monkeys’ desertion behavior as a function of reward prediction error during the result period:

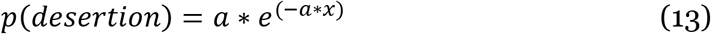

where *α* is the rate parameter. Paired t-tests were used to test for significance of any difference in desertion rate in the control and inactivation condition.

### DATA AND SOFTWARE AVAILABILITY

Data and software are available upon request to the Lead Contact.

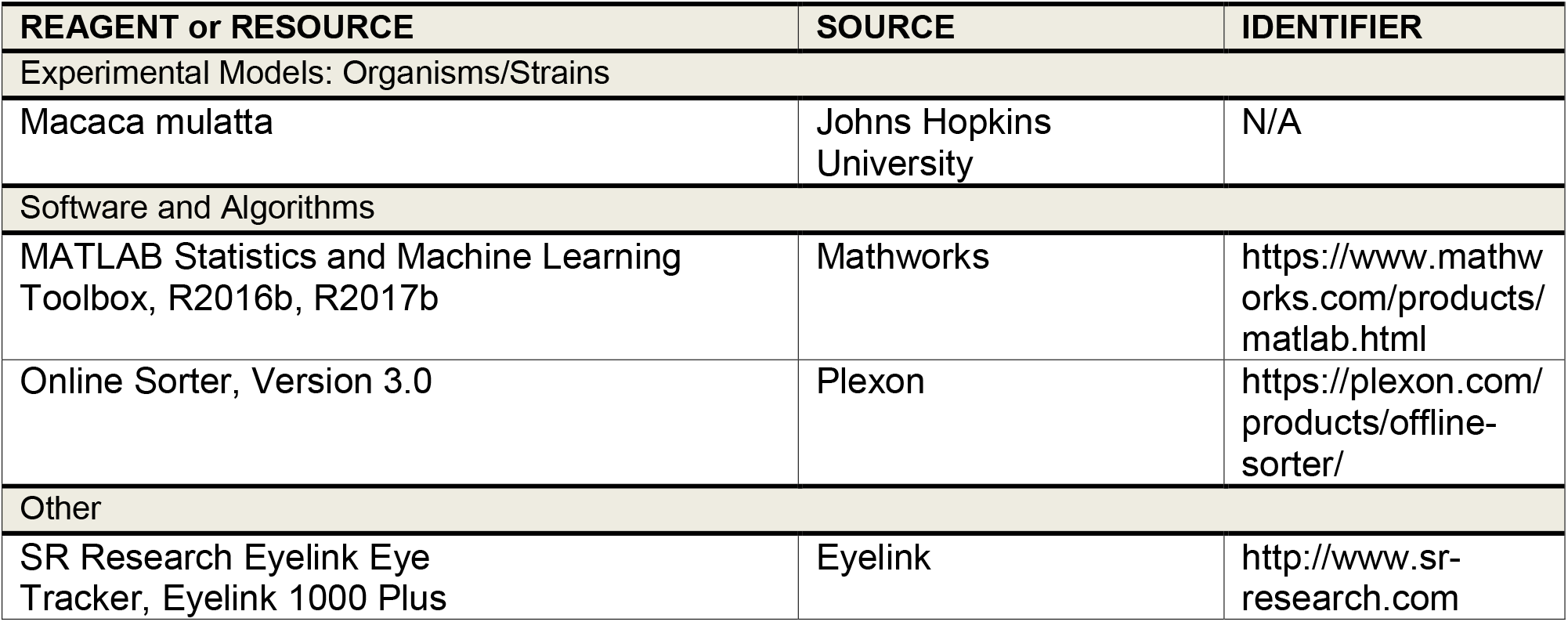

## Supplemental Data

**Figure S1.**
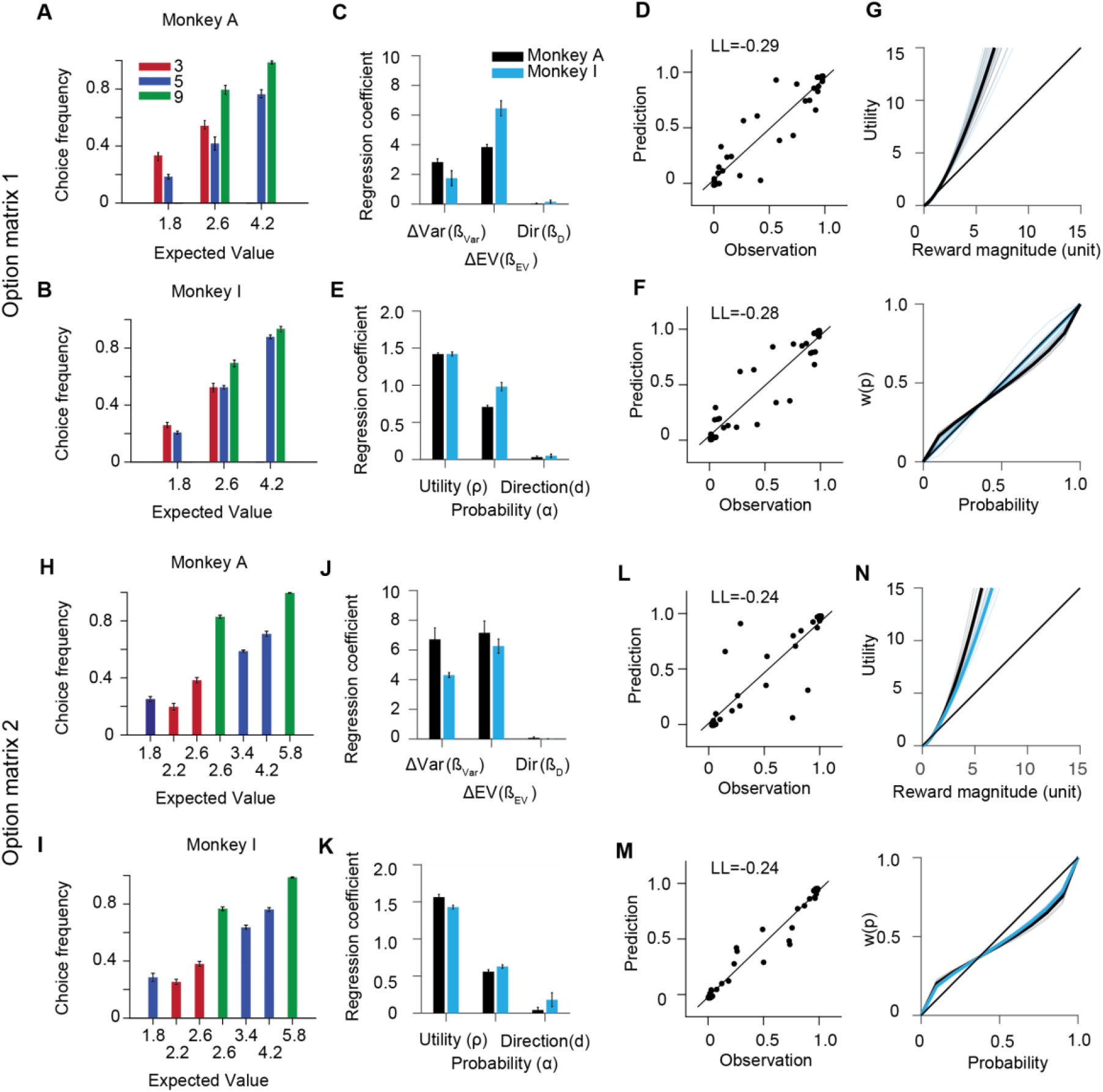
Choice patterns with both option matrices. Related to Figure 2. Quantitatively similar behavioral results were observed using both option matrix 1 and option matrix 2, when behavior was modeled using the risk-value and the prospect theory models. Across both option matrices, the monkeys behaved risk-seeking. (A-F) Option matrix 1. (A, B) The overall frequency of choosing a particular gamble option, when paired against all other options, for option matrix 1 for monkey A (A) and monkey I (B). The colors of the bars indicate the maximum reward amount of the gamble options (same as Figure 1B). (C) Regression coefficients of the risk-value model indicate preference for options with higher risk (ΔVar, β_Var_) and higher expected value (ΔEV, β_EV_) (Monkey A, black; Monkey I, blue). (D) Prediction accuracy of choice frequencies across the gamble options for the risk-value model. (E) Regression coefficients of the prospect theory model indicate a convex utility function (ρ) and an inverted S-shaped probability weighting function (α) (Monkey A, black; Monkey I, blue). (F) Prediction accuracy of choice frequencies across the gamble options for the prospect theory model. (G) The top panel shows the utility functions across all sessions (thick line) and for individual sessions (thin lines). The bottom panel shows the probability weighting functions across all sessions (thick line) and for individual sessions (thin lines). (Monkey A, thick black and thin grey lines; Monkey I, thick dark blue and thin light blue lines) (H-M) Option matrix 2: Same schema as for option matrix 1. Error bars denote s.e.m.

**Figure S2.**
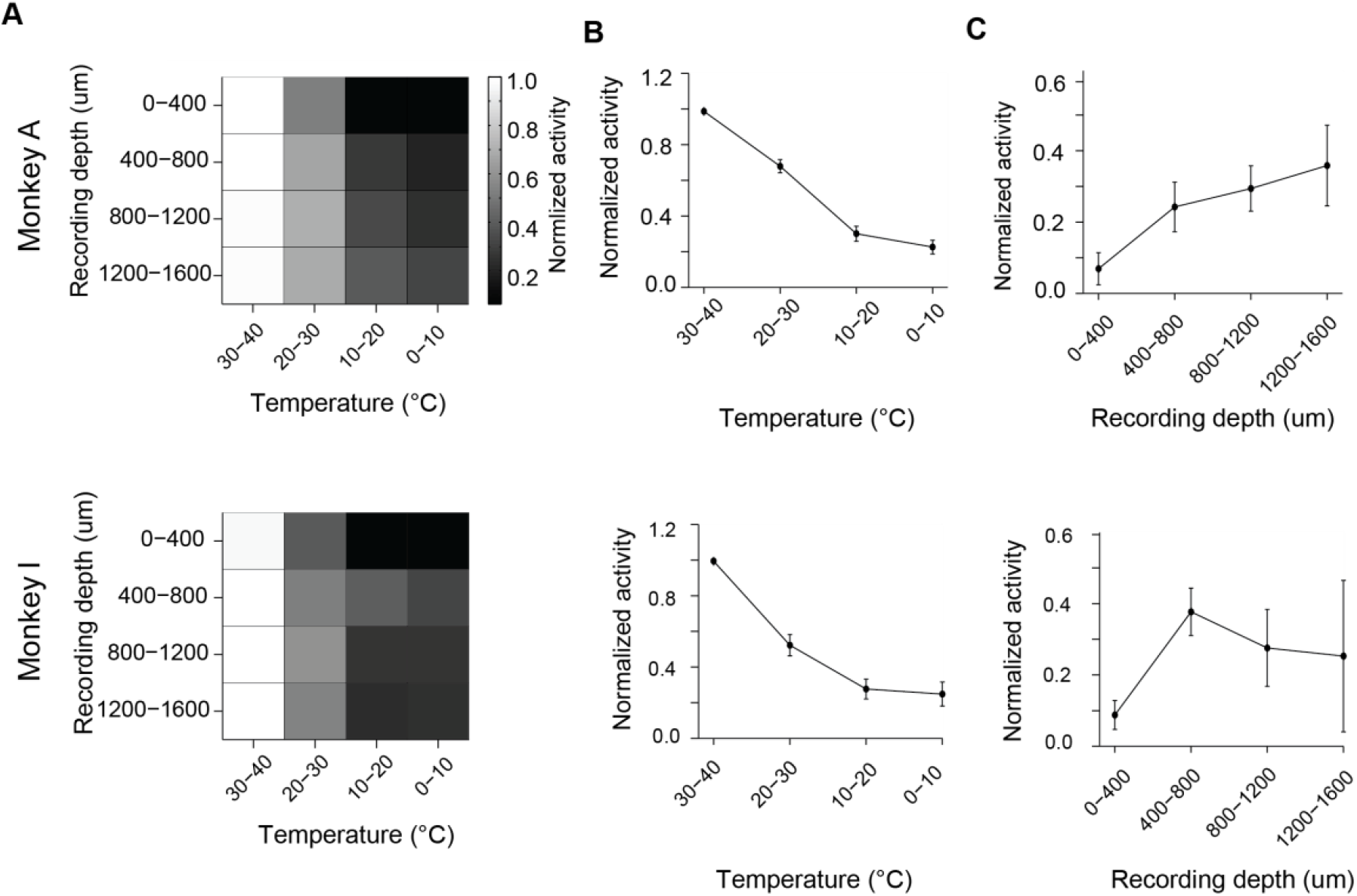
The effect of cooling inactivation on action potentials. Related to Figure 1 and STAR Methods. (A) The effect of cooling on SEF multi-unit spiking activity as a function of the temperature of the cooling probe and depth of recording for both Monkey A (top) and Monkey I (bottom). For each inactivation session, the average multi-unit firing rate during each trial was normalized to its maximum average firing rate across all trial conditions. The matrix shows the average normalized activity across all recordings for each monkey for a particular combination of temperature and recording depth. The brightness of the gray scale indicates activity levels. As the matrix indicates, maximum neuronal activity is seen during control temperature conditions in the absence of inactivation. The darkness of an element in the matrix therefore indicates the degree to which activity is reduced by inactivation. (B) The neuronal activity as a function of temperature averaged across all recording depths for both monkeys. (C) The neuronal activity as a function of recording depth averaged across all temperatures for both monkeys. b and c confirm the pattern seen in the matrix. The error bars represent s.e.m.. Lowering the temperature of the cooling probe leads to a reduction of neuronal activity. The activity reduction is more pronounced the closer the recording site is to the cooling probe above the cortical surface. Nevertheless, even at the deepest recording sites (1200-1600 *μ* m), the neuronal activity was still reduced by around 70% if temperatures were less than 10°C. Altogether, the distance over which the cooling affected the cortex was restricted to ≤2.5 mm. Accordingly, neighboring areas in the medial wall, such as pre-supplementary motor area and anterior cingulate cortex (ACC) should not be affected by the cooling.

**Figure S3.**
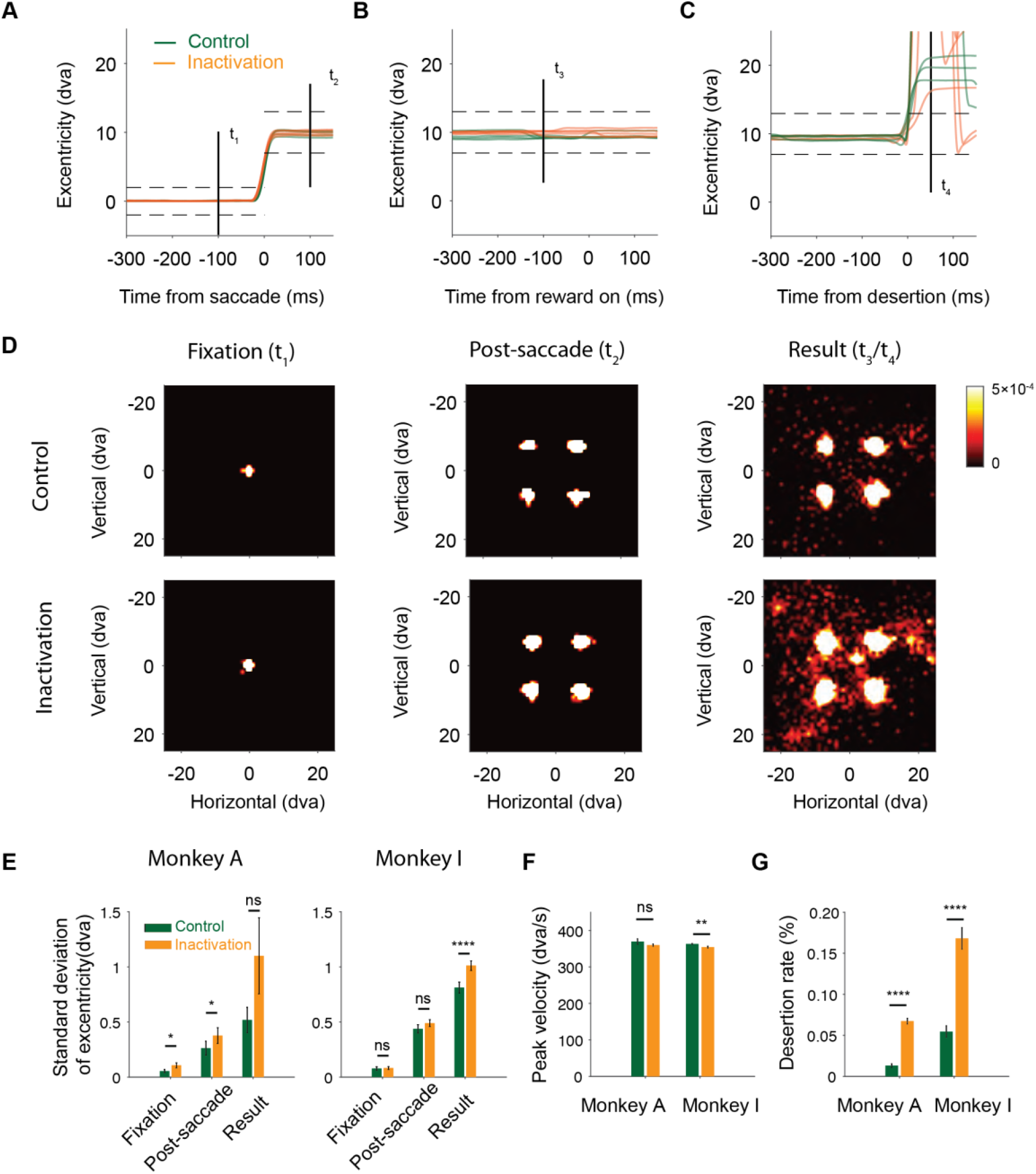
The effect of bilateral SEF inactivation on saccade and fixation metrics. Related to Figure 4. (A-C) Eye position traces during an example session in the control (n=5 trials, green) and inactivation (n=5 trials, orange) condition. The eye position traces were quantified by eccentricity (dva, degree of visual angle, radius from the center of the screen). The dashed lines indicate the positions of fixation or saccade windows. (A) The eye position traces during the choice period immediately before and after the saccades. The traces are aligned on the onset of the saccade, with which the monkey chooses the desired gamble option. (B) The eye position traces aligned on reward onset during the success trials, in which the monkey successfully finished the trials. The reward onset time is the same as the result cue turn-off time, and it is the time when fixation is no longer required. (C) The eye position traces aligned on desertion onset in the desertion trials during result period. In these trials, the monkeys fail to hold fixation by making a saccade outside of the fixation window after the gamble results were revealed. (D) Eye position density estimates during the fixation (t1, left), post-saccade (t2, middle), and result (t3 and t4 combined, right) periods when fixation was required to finish the trial. The fixation period shows the eye position distribution during the fixation period (t1=100ms before saccade onset, see a). The distribution during post-saccade period shows the scatter of the fixations shortly after saccade to the choice option (t2=100ms after saccade onset, see a). The distribution during result period (t3=100ms before the reward was delivered or 50ms after the trials were deserted, see b and c) shows the eye positions of the monkeys during the time period after the gamble results were revealed. During this period, fixations were required for the reward delivery. (E) Standard deviations of eccentricities of fixations during fixation (t1), post-saccade (t2), and result periods in the success trials (t3) in control (green) and inactivation (orange) conditions. Inactivation significantly increase of fixation scatters during fixation and post-saccade periods for Monkey A, and during result periods for Monkey I. Importantly, the significant increased variances in eye position during the different periods were less than 0.2 dva. For comparison, the size of the fixation window was 4 dva, and the size of the saccade target window was 6 dva. (F) Peak velocities of saccades during choice period show no significant differences between control and inactivation condition for monkey A. They are slightly slower (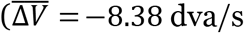, paired t-test, *p*=0.01) for Monkey I. (G) Desertion rates during the result period. The desertion trials are defined as the trials in which the monkeys made active saccades outside of the fixation window after the gamble results were revealed. The desertion rates are significantly higher for both monkeys (Monkey A: 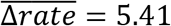, paired t-test, *p* = 1.00 × 10^-8^; Monkey I: 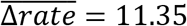, paired t-test, *p* = 1.24 × 10^-7^). The error bars represent s.e.m; paired t-test; ns, nonsignificant; *, *p* <0.05; **, *p* <10^-2^; ***, *p* <10^-3^; ****, *p* <10^-4^.

**Figure S4.**
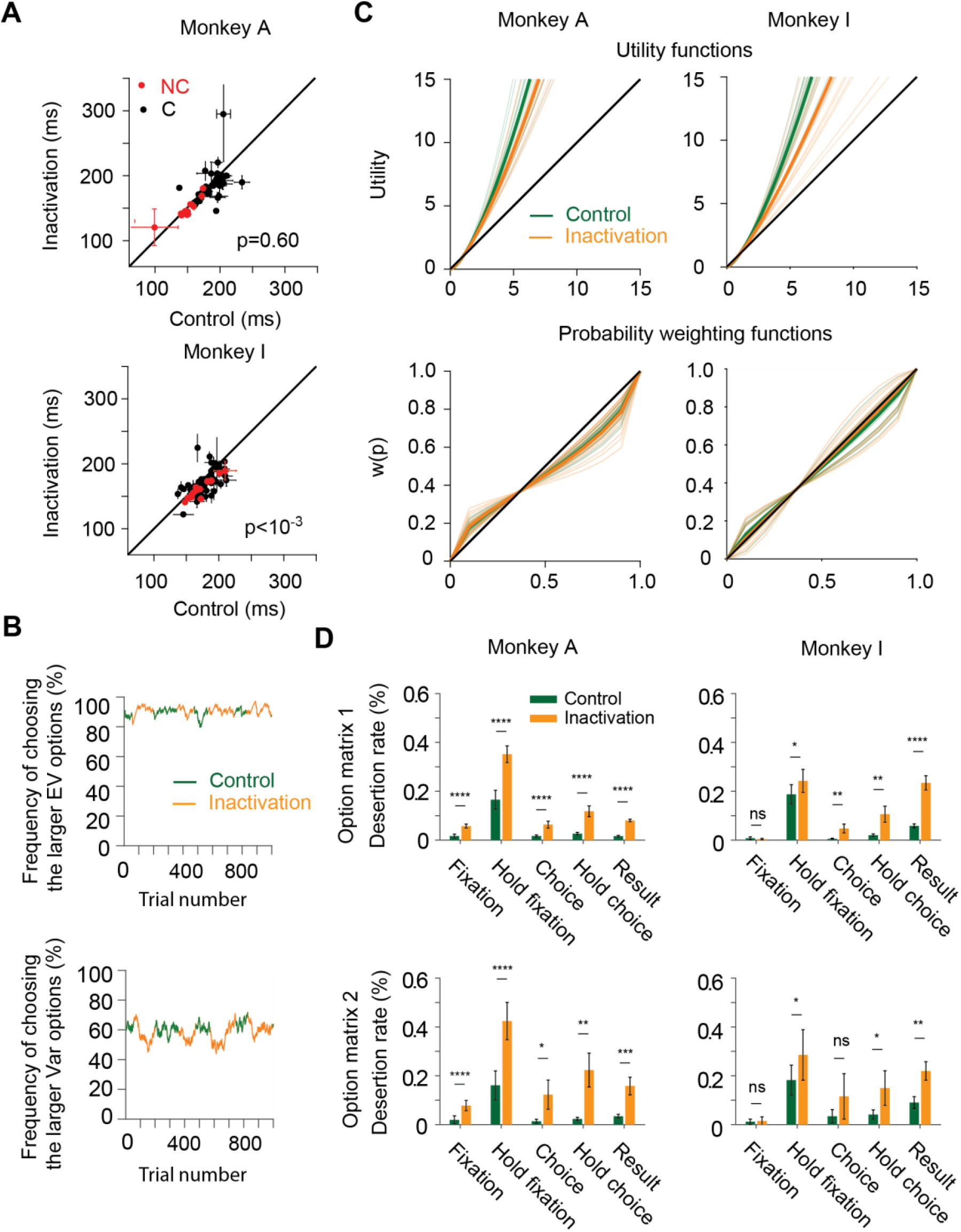
The effect of bilateral SEF inactivation on reaction time, risk preference and trial dissertation rate. Related to Figure 3 and 4. (A) The effect of bilateral SEF inactivation on reaction times for no-choice trials (14 trial types) and choice trials (42 trial types) for monkey A (top) and monkey I (bottom). There is no significant change of reaction times for monkey A in both no-choice (*p* = 0.23) and choice condition (*p* = 0.78). There is a small but significant reduction of reaction times for monkey I in both no-choice (*p* = 1.08 × 10^-4^) and choice condition (*p* = 5.30 × 10^-3^) (see also table S3). (B) An example session shows the trial-by-trial change of choice frequency for higher EV options (top) and higher Var options (bottom) during control (green) and inactivation (orange). (C) Reduction of risk-seeking estimated by the prospect theory model. The corresponding utility functions (top) and probability weighing functions (bottom) are shown during the control (green) and inactivation (orange) condition for both monkey A (left) and I (right). (D) Bilateral SEF inactivation increased the trial quitting rate of trial in almost all epochs of the task. The panels compare dissertation rates during 5 different trial periods in both monkeys across both option matrices in the control (green) and inactivation (orange) condition. Specifically, trial dissertation in each epoch is defined as: 1) Fixation: failing to saccade into the fixation window within a 1s time window following the onset of the fixation cue. 2) Hold fixation: breaking fixation during the 500-1000ms period when only the fixation spot was on the screen, before the targets appear. 3) Choice: failing to choose a target by making a saccade into one of the target windows within a 1 s time window following target onset (i.e., the decision-period). 4) Hold Choice: breaking fixation during the 500-600 ms period following the choice, before the gamble results were revealed. 5) Result: breaking fixation during the 300–600 ms period after the result was revealed and before the reward is delivered. Error bars denote s.e.m.; paired t-test, ns, non-significant; *, *p* <0.05; **, *p* <10^-2^; ***, *p* <10-3; ****, *p* <10^-4^.

**Table S1.** Gamble options used in both option matrices. Related to Figure 1B

| Options |  | 1 | 2 | 3 | 4 | 5 | 6 | 7 |
| --- | --- | --- | --- | --- | --- | --- | --- | --- |
| Option Matrix 1 | Maximum reward | 3 | 3 | 5 | 5 | 5 | 7 | 7 |
|  | Probability of winning | 40% | 80% | 20% | 40% | 80% | 20% | 40% |
|  | Minimum reward | 1 | 1 | 1 | 1 | 1 | 1 | 1 |
|  | Probability of losing | 60% | 20% | 80% | 60% | 20% | 80% | 60% |
|  | Expected value | 1.8 | 2.6 | 1.8 | 2.6 | 4.2 | 2.6 | 4.2 |
|  | Variance | 0.98 | 0.80 | 1.60 | 1.96 | 1.60 | 3.20 | 3.92 |
|  | Coefficient of variance | 0.54 | 0.31 | 0.89 | 0.75 | 0.38 | 1.23 | 0.93 |
| Option Matrix 2 | Maximum reward | 3 | 3 | 5 | 5 | 5 | 7 | 7 |
|  | Probability of winning | 60% | 80% | 20% | 60% | 80% | 20% | 60% |
|  | Minimum reward | 1 | 1 | 1 | 1 | 1 | 1 | 1 |
|  | Probability of losing | 40% | 20% | 80% | 40% | 20% | 80% | 40% |
|  | Expected value | 2.2 | 2.6 | 1.8 | 3.4 | 4.2 | 2.6 | 5.8 |
|  | Variance | 0.98 | 0.80 | 1.60 | 1.96 | 1.60 | 3.20 | 3.92 |
|  | Coefficient of variance | 0.45 | 0.31 | 0.89 | 0.58 | 0.38 | 1.23 | 0.68 |

**Table S2.**
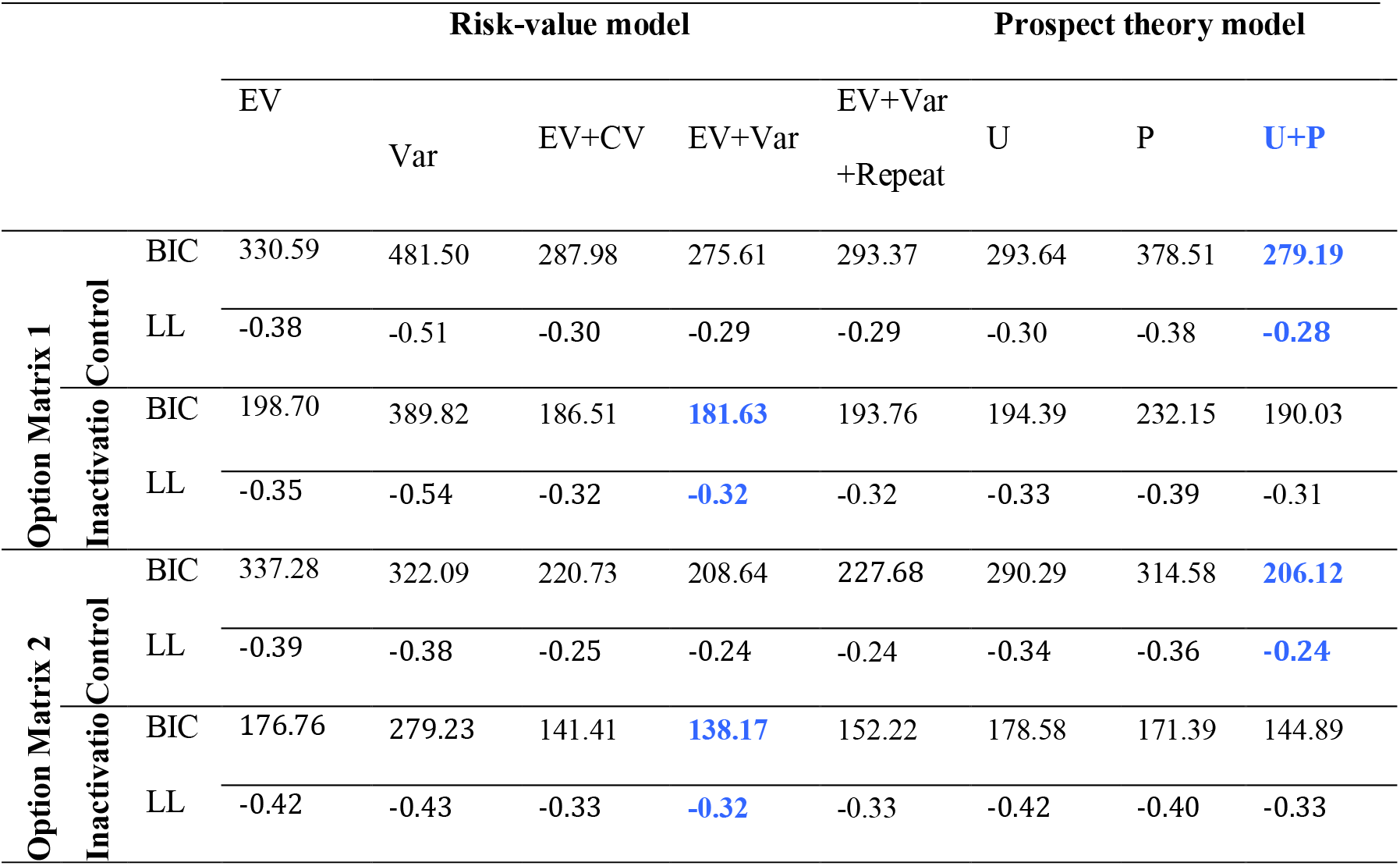
Bayesian information criterion (BIC) and log-likelihood (LL) for different risk models. Related to Figure 2.

BIC values are computed averaging across session across two monkeys. LL values are computed using cross validation for monkeys (see Methods). The prospect theory models with both nonlinear utility function and probability weighting function are the best models (blue) with lowest BIC values for all control conditions.

**Table S3.**
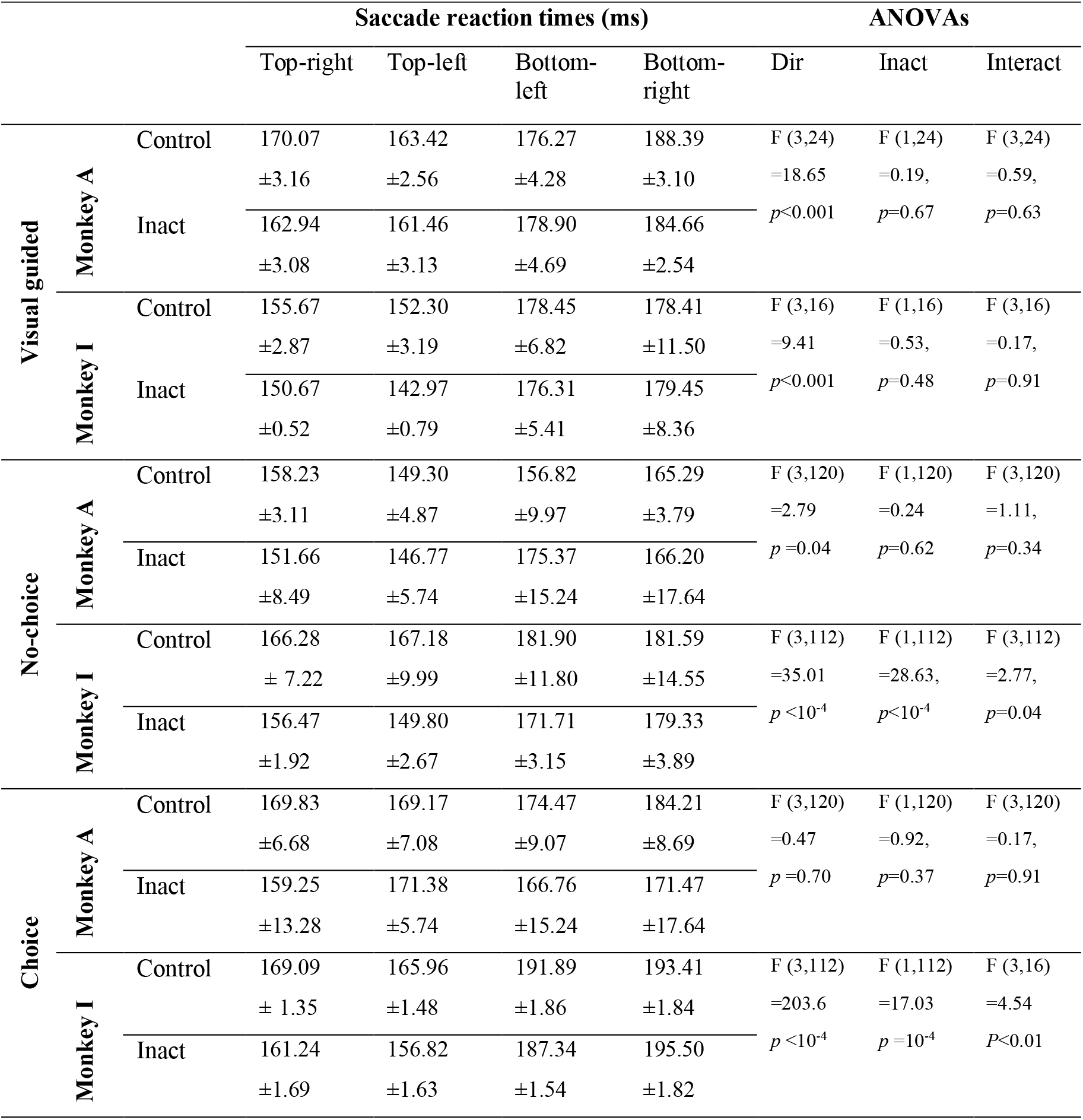
SEF inactivation does not have significant/consistent effect on saccade reaction times in both visual guided saccade task and gamble task. Related to Figure 3.

Saccade reaction times columns represent the monkeys’ saccade reaction time ± s.e.m. for each direction during inactivation (Inact) and control (Control) conditions. ANOVAs test columns represent the *p* value in ANOVAs test for a direction effect (Dir), an inactivation effect (Inact), and an interaction effect (Interact) between direction and inactivation on reaction time during the visual guided saccade task. The results show that both monkeys showed a directional bias in their reaction time, but no consistent inactivation effect.

